# Re-evaluation of SNP heritability in complex human traits

**DOI:** 10.1101/074310

**Authors:** Doug Speed, Na Cai, The UCLEB Consortium, Michael R. Johnson, Sergey Nejentsev, David J Balding

**Affiliations:** UCL Genetics Institute, University College London, United Kingdom; Wellcome Trust Sanger Institute, Wellcome Genome Campus, Hinxton, Cambridge, United Kingdom; European Molecular Biology Laboratory, European Bioinformatics Institute (EMBL-EBI), Wellcome Genome Campus, Hinxton, Cambridge, United Kingdom; Division of Brain Science, Imperial College London, United Kingdom; Department of Medicine, University of Cambridge, United Kingdom; Centre for Systems Genomics, School of BioSciences and School of Mathematics & Statistics, University of Melbourne, Australia

## Abstract

SNP heritability, the proportion of phenotypic variance explained by SNPs, has been reported for many hundreds of traits. Its estimation requires strong prior assumptions about the distribution of heritability across the genome, but the assumptions in current use have not been thoroughly tested. By analyzing imputed data for a large number of human traits, we empirically derive a model that more accurately describes how heritability varies with minor allele frequency, linkage disequilibrium and genotype certainty. Across 19 traits, our improved model leads to estimates of common SNP heritability on average 43% (SD 3) higher than those obtained from the widely-used software GCTA, and 25% (SD 2) higher than those from the recently-proposed extension GCTA-LDMS. Previously, DNaseI hypersensitivity sites were reported to explain 79% of SNP heritability; using our improved heritability model their estimated contribution is only 24%.

---

The SNP heritability (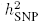) of a trait is the fraction of phenotypic variance explained by additive contributions from SNPs.^1^ Accurate estimates of 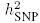 are central to resolving the missing heritability debate, indicate the potential utility of SNP-based prediction and help design future genome-wide association studies (GWAS).^2,3^ Whereas techniques for estimating (total) heritability have existed for decades,^4,5^ the first method for estimating 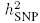 was proposed only in 2010,^1^ but has since been applied to many hundreds of traits. Extensions of this method are now being used to partition heritability across chromosomes, biological pathways and by SNP function, and to calculate the genetic correlation between pairs of traits.^6–8^

As the number of SNPs in a GWAS is usually much larger than the number of individuals, estimation of 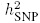 requires steps to avoid over-fitting. Most reported estimates of 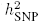 are based on assigning the same Gaussian prior distribution to each SNP effect size, in a way which implies that all SNPs are expected to contribute equal heritability.^1,9^ By examining a large collection of real datasets, we derive approximate relationships between the expected heritability of a SNP and minor allele frequency (MAF), levels of linkage disequilibrium (LD) with other SNPs and genotype certainty. This provides us with an improved model for heritability estimation and a better understanding of the genetic architecture of complex traits.

## Results

When estimating 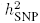, the “LDAK Model” assumes

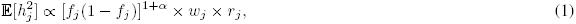

where 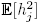 is the expected heritability contribution of SNP *j* and *f_j_* is its (observed) MAF. The parameter α determines the assumed relationship between heritability and MAF. In human genetics it is commonly assumed that heritability does not depend on MAF, which is achieved by setting α = –1, however, we consider alternative relationships. The SNP weights *w*_1_,…, *w_m_* are computed based on local levels of LD;^9^ *w_j_* tends to be higher for SNPs in regions of low LD, and thus the LDAK Model assumes that these SNPs contribute more than those in high-LD regions. Finally, *r_j_* ∈ [0, 1] is an information score measuring genotype certainty; the LDAK Model expects that higher-quality SNPs contribute more than lower-quality ones. *r_j_* is defined in Online Methods, where we also explain how (1) arises by assuming a genome-wide random regression in which SNP effect sizes are assigned Gaussian distributions.

The “GCTA Model” is obtained from (1) by setting *w_j_* = 1 and *r_j_* = 1, and thus assumes that expected heritability does not vary with either LD or genotype certainty. To date, most reported estimates of 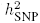 have used the GCTA Model with α = –1, which corresponds to the assumption that 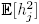 is constant, and so the expected contribution of a SNP set depends only on the number of SNPs it contains.^1^ To appreciate the major difference between the GCTA and LDAK Models, consider a region containing two SNPs: under the GCTA Model, the expected heritability of these two SNPs is the same irrespective of the LD between them, whereas under the LDAK Model, two SNPs in perfect LD are expected to contribute only half the heritability of two SNPs showing no LD. See Figure 1 for a more detailed example.

**Figure 1:**
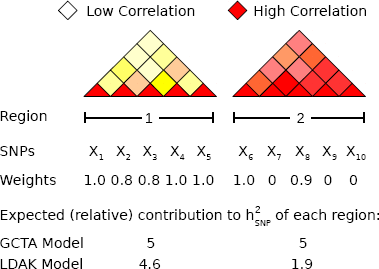
Comparison of the GCTA and LDAK Models. Region 1 contains five SNPs in low LD (lighter colors indicate weaker pairwise correlations). Each SNP contributes unique genetic variation, reflected by SNP weights close to one. Region 2 contains five SNPs in high LD (strong correlations). The total genetic variation tagged by the region is effectively captured by two of the SNPs, and so the others receive zero weight. Under the GCTA Model, the regions are expected to contribute heritability proportional to their numbers of SNPs, here equal. Under the LDAK Model, they are expected to contribute proportional to their sums of SNP weights, here in the ratio 4.6:1.9. Note that the expected heritability can also depend on the allele frequencies and genotype certainty of the SNPs, but for simplicity, these factors are ignored here.

An alternative method for estimating 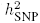 is LDSC (LD Score Regression).^10^ The LDSC Model expects that each SNP contributes equal heritability,^10,11^ and therefore closely resembles the GCTA Model with α = –1. When applied to the same dataset, estimates from LDSC will typically have standard error 25-100% higher than those from GCTA;^11^ this is partly because the LDSC Model includes an extra parameter, designed to capture confounding biases, and partly because LDSC estimates are moment-based, whereas GCTA (like LDAK) uses restricted maximum likelihood (REML).^12,13^ However, as LDSC requires only summary statistics (i.e., *p*-values from single-SNP analysis), it can be used on much larger datasets than GCTA and LDAK, which need raw genotype data, and can be applied to results from large-scale meta-analyses.^10^

### SNP partitioning

(1) can be generalized by dividing SNPs into tranches across which the constant of proportionality is allowed to vary (so 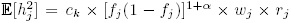 SNPs in Tranche *k*). This is known as SNP partitioning.^6^ Two examples are GCTA-MS^14^ and GCTA-LDMS:^15^ when applied to common SNPs (MAF > 0.01), GCTA-MS divides the genome into five tranches based on MAF, using the boundaries 0.1, 0.2, 0.3 and 0.4, while GCTA-LDMS first divides SNPs into four tranches based on local average LD Score,^10^ then divides each of these into five based on MAF, resulting in a total of 20 tranches. In general, we prefer to avoid SNP partitioning when estimating 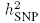, because it introduces (often arbitrary) discontinuities in the model assumptions and can cause convergence problems. However, we show below that partitioning based on MAF enables reliable estimation of 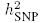 when rare SNPs (MAF < 0.01) are included. Additionally, SNP partitioning provides a way to visually assess the fit of different heritability models; it allows us to estimate average 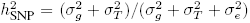 for different SNP tranches, which can then be compared to the values predicted under different assumptions.

### Datasets

In total, we analyze data for 42 traits. Table 1 describes the 19 “GWAS traits” (17 case-control, 2 quantitative). For these, individuals were genotyped using either genome-wide Illumina or Affymetrix arrays (typically 500 K to 1.2 M SNPs). We additionally examine data from eight cohorts of the UCLEB consortium,^24^ which comprise about 14 000 individuals genotyped using the Metabochip;^25^ (a relatively sparse array of 200 K SNPs selected based on previous GWAS) and recorded for a wide range of clinical phenotypes. From these, we consider 23 quantitative phenotypes (average sample size 8 200), which can loosely be divided into anthropomorphic (height, weight, BMI and waist circumference), physiological (lung capacity and blood pressure), cardiac (e.g., PR and QT intervals), metabolic (glucose, insulin and lipid levels) and blood chemistry (e.g., fibrinogen, Interleukin 6 and haemoglobin levels). In general, our quality control is extremely strict; after imputing using IMPUTE2^26^ and the 1000 Genome Phase 3 (2014) reference panel,^27^ we retain only autosomal SNPs with MAF > 0.01 and information score *r_j_* > 0.99. We only relax quality control when, using the UCLEB data, we explicitly examine the consequences of including lower-quality and rare SNPs.

**Table 1:**
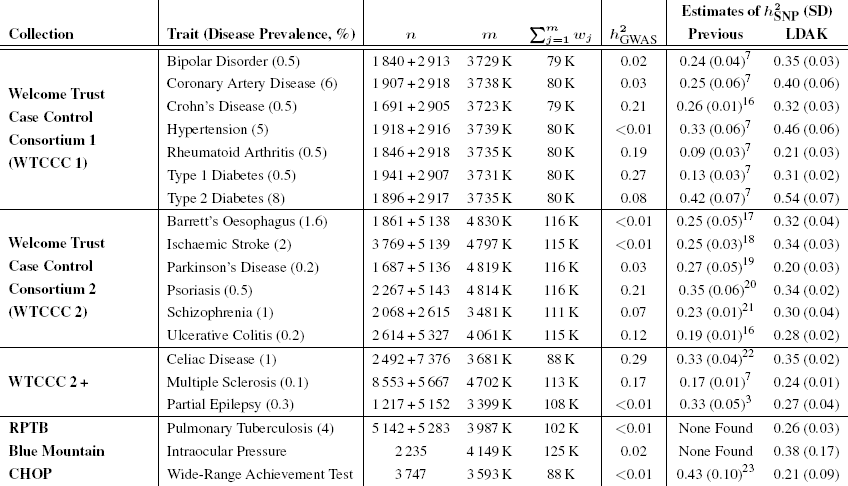
**Properties of datasets and estimates of 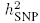**.*n* = sample size (cases + controls), *m* = number of SNPs, 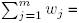 sum of SNP weights which can be interpreted as an effective number of independent SNPs. All values are post quality control; values for *m* and ∑*w_j_* are rounded to the nearest K (thousand). For UCLEB, *m* and ∑*w_j_* refer to our main analysis, which considers only high-quality, common SNPs. The final column provides our best estimates of 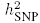 from common SNPs, computed using LDAK with α = –0.25 (see main text for explanation of α). For comparison, we include previously published estimates of 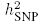 (note that the previous analyses for rheumatoid arthritis, type 1 diabetes and multiple sclerosis excluded major histocompatibility SNPs, which we estimate contribute 0.07, 0.20 and 0.05, respectively), as well as 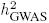, the proportion of phenotypic variance explained by SNPs reported as GWAS significant (*P* < 5 × 10^−8^). For disease traits, estimates of 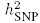 and 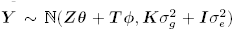 have been converted to the liability scale assuming the stated prevalence.

Further details of our methods and datasets are provided in Online Methods. In particular, we explain how when estimating 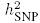 we give special consideration to highly-associated SNPs, which we define as those with *P* < 10^−20^ from single-SNP analysis, and how for the UCLEB data, we confirm that genotyping errors do not correlate with phenotype (which is important for the analyses where we include lower-quality SNPs).

### Relationship between heritability and MAF

Varying the value of α in (1) changes the assumed relationship between heritability and MAF; three example relationships are shown in Figure 2a. To determine suitable α, we analyze each of the 42 traits using seven values: –1.25, –1, –0.75, –0.5, –0.25, 0 and 0.25, seeing which lead to best model fit (highest likelihood). Full results are provided in Supplementary Figure 1 and Supplementary Table 2. First, to remove any confounding due to LD, we use only a pruned subset of SNPs (with *w_j_* = 1); next, we repeat without LD pruning (the results for the GWAS traits are shown in Figure 2b); finally, for the UCLEB traits, we repeat including lower-quality and rare SNPs. We find that model fit is typically highest for –0.5 ≤ α ≤ 0, whereas the most widely-used value, α = –1, reuslts in sub-optimal fit. On the basis that it performs consistently well across different traits and SNP filterings, we recommend that α = –0.25 becomes the default. This value implies that expected heritability declines with MAF; this is seen in Figure 2a which reports, averaged across the 19 GWAS traits, the (weight-adjusted) per-SNP heritability for low- and high-MAF SNPs (see Supplementary Figure 2 for further details).

**Figure 2:**
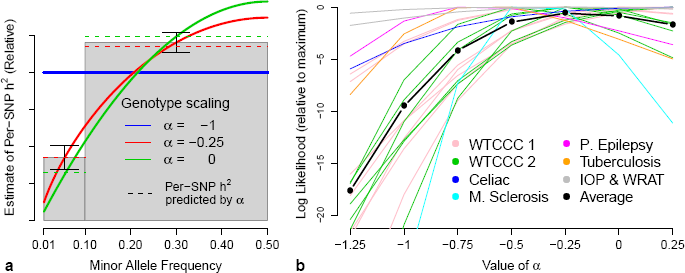
(a) Relationship between heritability and MAF. The parameter α specifies the assumed relationship between heritability and MAF: in human genetics, α = –1 is typically used (solid blue line), while in animal and plant genetics, α = 0 is more common (green); we instead found α = –0.25 (red) provides a better fit to real data. The gray bars report (relative) estimates of the per-SNP heritability for MAF < 0.1 and MAF>0.1 SNPs, averaged across the 19 GWAS traits (vertical lines provide 95% confidence intervals); the dashed lines indicate the per-SNP heritability predicted by each α. **(b) Determining best-fitting** α **for the GWAS traits**. We compare α based on likelihood; higher likelihood indicates better-fitting α. Lines report log likelihoods from LDAK for seven values of α, relative to the highest observed. Line colors indicate the seven trait categories, while the black line reports averages.

While α = –0.25 provides the best fit overall, for individual traits, optimal α may differ, and therefore we investigate sensitivity of 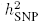 estimates to the value of α. Full results are provided in Supplementary Figures 3, 4 & 5, while Figure 6a provides a summary for the UCLEB traits. When analyzing only common SNPs, we find that changes in α have little impact on 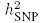. For example, across the 23 UCLEB traits, estimates from high-quality common SNPs using α = –0.25 are on average only 5% (SD 4) lower than those using α = –1, and 4% (SD 4) higher than those using α = 0. However, this is no longer the case when rare SNPs are included in the analysis: for example, when the MAF threshold is reduced to 0.0005, estimates using α = –0.25 are on average 18% (SD 4) lower than those using α = –1 and 30% (SD 6) higher than those from α = 0. Therefore, when including rare SNPs, we guard against misspecification of α by partitioning based on MAF (with boundaries at 0.001, 0.0025, 0.01 and 0.1); we find that this provides stable estimates of 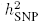 and also allows estimation of the relative contributions of rare and common variants (Figure 6a and Supplementary Figure 6).

**Figure 3:**
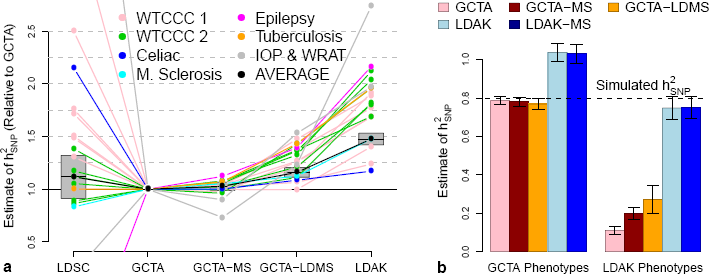
(a) Relative estimates of 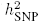 for the GWAS traits. 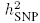 estimates from LDSC, GCTA-MS (SNPs partitioned by MAF), GCTA-LDMS (SNPs partitioned by LD and MAF) and LDAK are reported relative to those from GCTA. For versions of GCTA and LDAK, we use α = –0.25 (see main text for explanation of α). Line colors indicate the seven trait categories; the black line reports the (inverse variance weighted) averages, with gray boxes providing 95% confidence intervals for these averages. Numerical values are provided in Supplementary Table 3. **(b) Simulation studies can be misleading**. Phenotypes are simulated with 1000 causal SNPs and 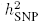 = 0.8 (black horizontal line), then analyzed using GCTA, GCTA-MS, GCTA-LDMS, LDAK and LDAK-MS (LDAK with SNPs partitioned by MAF). Bars report average 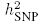 across 200 simulated phenotypes (vertical lines provide 95% confidence intervals). **Left**: copying the study of Yang *et al.*,^1^ causal SNP effect sizes are sampled from ℕ(0, 1), similar to the GCTA Model. **Right**: causal SNP effect sizes are sampled from ℕ(0, *w_j_*), similar to the LDAK Model.

**Figure 4:**
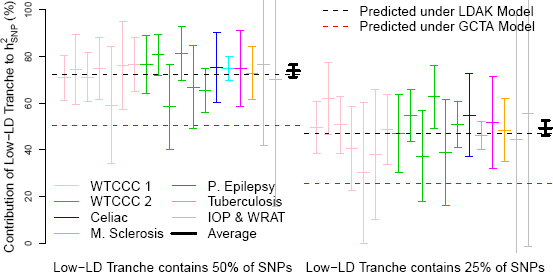
Comparing the GCTA and LDAK Models for the GWAS traits. We partition SNPs into low-or high-LD, with the low-LD tranche containing either 50% (left) or 25% (right) of SNPs. For each partition, the horizontal red and black lines indicate the predicted contribution of the low-LD tranche to 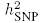 under the GCTA and LDAK Models, respectively. Vertical lines provide point estimates and 95% confidence intervals for the contribution of the low-LD tranche to 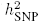, estimated assuming the GCTA Model. Line colors indicate the seven trait categories, while the black lines provide the (inverse variance weighted) averages.

**Figure 5:**
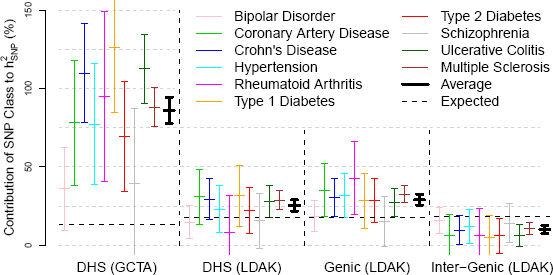
Enrichment of SNP Classes. Block 1 reports the contributions to 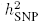 of DNaseI hypersensitivity sites (DHS), estimated under the GCTA Model with α = –1 (see main text for explanation of α). The vertical lines provide point estimates and 95% confidence intervals for each trait, and for the (inverse variance weighted) average; for 3 of the traits, the point estimate is above 100%, as was also the case for Gusev *et al.*^7^ Block 2 repeats this analysis, but now assuming the LDAK Model with α = –0.25. Blocks 3 & 4 estimate the contribution of “genic SNPs” (those inside or within 2 kb of an exon) and “inter-genic SNPs” (further than 125 kb from an exon), again assuming the LDAK Model with α = –0.25. To assess enrichment, estimated contributions are compared to those expected under the GCTA or LDAK Model, as appropriate (horizontal lines).

**Figure 6:**
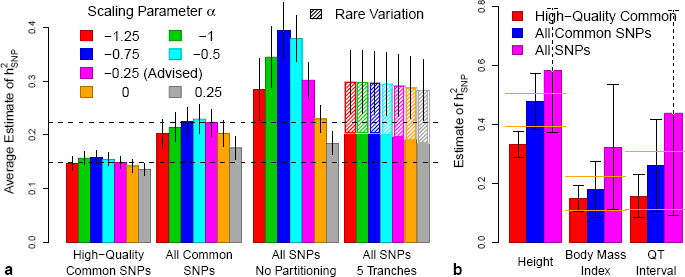
Varying quality control for the UCLEB traits. We consider three SNP filterings: 353 K high-quality common SNPs (information score > 0.99, MAF > 0.01), 8.8 M common SNPs (MAF > 0.01) and all 17.3 M SNPs (MAF > 0.0005). (**a**) Blocks indicate SNP filtering; bars report (inverse variance weighted) average estimates of 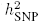 using LDAK (vertical lines provide 95% confidence intervals). Bar color indicates the value of α used. For Blocks 1, 2 & 3, 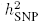 is estimated using the non-partitioned model. For Block 4, SNPs are partitioned by MAF; we find this is necessary when rare SNPs are included, and also allows estimation of the contribution of MAF < 0.01 SNPs (hatched areas). (**b**) bars report our final estimates of 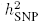 for height, body mass index and QT interval, the three traits for which common SNP heritability has been previously estimated with reasonable precision^6^ (orange lines mark the 95% confidence intervals from these previous studies). Bar colors now indicate SNP filtering; all estimates are based on α = –0.25, using either a non-partitioned model (red and blue bars) or with SNPs partitioned by MAF (purple bars).

### Relationship between heritability and LD

The LDAK Model assumes that heritability varies according to local levels of LD, whereas the GCTA Model assumes that heritability is independent of LD. First we demonstrate that choice of model matters when estimating 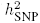. For the GWAS traits, Figure 3a reports relative estimates of 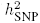 from GCTA, GCTA-MS, GCTA-LDMS and LDAK (all using α = –0.25); see Supplementary Figure 7 for an extended version. We find that estimates based on the LDAK Model are on average 48% (SD 3) higher than estimates based on the GCTA Model. For the UCLEB traits, estimates from LDAK are on average 88% (SD 7) higher than those from GCTA (Supplementary Fig. 8). Figure 3a also includes results from LDSC, run as described in the original publication^10^ (see Supplementary Table 3 for numerical values). Estimates from LDSC are not significantly different to those from GCTA, which is to be expected considering that GCTA and LDSC assume the same relationship between heritability and LD. In Supplementary Figure 9 we consider alternative versions of LDSC (e.g., varying how LD Scores are computed, forcing the intercept term to be zero and excluding highly-associated SNPs). While changing settings can have a large impact, in all cases the average estimate of 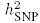 from LDSC remains substantially below that from LDAK.

A recent article which asserted that GCTA estimates 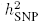 more accurately than LDAK, based this claim on a simulation study in which causal SNPs were assigned effect sizes from the same Gaussian distribution, irrespective of LD.^6^ This resembles the GCTA Model but not the LDAK Model, and so it is no surprise that GCTA performed better. Figure 3b shows that if instead effect size variances had been scaled by SNP weights, and so vary with LD similar to the LDAK Model, then the study would have found LDAK to be superior to GCTA. Thus using simulations to compare different heritability models is problematic, because the conclusions will depend on the assumptions used when generating phenotypes. See Supplementary Figure 10 for a full reanalysis of the reported simulation study and Supplementary Figure 11 for further simulations.

Rather than using simulations, we compare LDAK and GCTA empirically. Supplementary Table 4 shows that when α = –0.25, assuming the LDAK Model leads to higher likelihood than assuming the GCTA Model for all 19 GWAS traits and for 17 of the 23 UCLEB traits (if we instead use α = –1, likelihood is higher under the LDAK Model for 31 of the 42 traits). To visually demonstrate the superior fit of the LDAK Model, we partition SNPs into low- and high-LD (for this, we rank SNPs according to the average LD Score^10^ of non-overlapping 100 kb segments, the metric used by GCTA-LDMS^15^). First, we partition so that the two tranches contain an equal number of SNPs. The left half of Figure 4b reports, for each of the GWAS traits, the contribution of the low-LD tranche, estimated using the GCTA Model (with α = –0.25). Under the GCTA Model, the low-LD tranche is expected to contribute 50% of 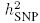; under the LDAK Model, it is expected to contribute 72% of 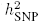. We see that the estimated contribution of the low-LD tranche is consistent with the GCTA Model (95% confidence interval includes 50%) for only 5 of the 19 traits, whereas it is consistent with the LDAK Model (confidence interval includes 72%) for 18. Next we partition so that the low-LD tranche contains a quarter of the SNPs; now the low-LD tranche is predicted to contribute 26% of 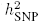 under the GCTA Model, but 47% of 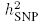 under the LDAK Model. The right half of Figure 4b shows that its estimated contribution is consistent with the GCTA Model for only 7 of the 19 traits, but again consistent with the LDAK Model for 18. Additional results are provided in Supplementary Figure 12; these show that regardless of whether we estimate heritabilities using LDAK (rather than GCTA), whether we use α = –1 (instead of α = –0.25) or whether we analyze the UCLEB traits, it remains the case that the LDAK Model better predicts the heritability contribution of each tranche than the GCTA Model.

### Relationship between heritability and genotype certainty

The LDAK Model assumes that SNP heritability contributions vary with genotype certainty (measured by the information score *r_j_*). So far, our analyses have used only very high-quality SNPs (*r_j_* > 0.99), so this assumption has been redundant. Now we also include lower-quality common SNPs; we focus on the UCLEB traits, as for these we were able to test for correlation between genotyping errors and phenotype (Supplementary Fig. 13). Supplementary Table 5 compares model fit with and without allowance for genotype certainty; it shows that including *r_j_* in the heritability model tends to provide a modest improvement in model fit, resulting in a higher likelihood for 18 out of 23 traits.

### Estimates of 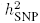 for the GWAS traits

Table 1 presents our final estimates of 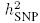 for the 19 GWAS traits, obtained using the LDAK Model (with α = –0.25). For comparison, we include previously-reported estimates of 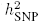, as well as the proportion of phenotypic variance explained by SNPs reported as genome-wide significant (see Supplementary Table 6). For the disease traits, estimates are on the liability scale, obtained by scaling according to the observed case-control ratio and (assumed) trait prevalence.^28,29^ We are unable to find previous estimates of 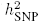 for tuberculosis or intraocular pressure, indicating that for these two traits, we are the first to establish that common SNPs contribute sizable heritability. Extended results are provided in Supplementary Table 7. These show that our final estimates of 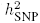 are on average 43% (SD 3) and 25% (SD 2) higher than, respectively, those obtained using the original versions (i.e., with α = –1) of GCTA^30^ and GCTA-LDMS.^15^

### Role of DNaseI hypersensitivity sites (DHS)

Gusev *et al.*^7^ used SNP partitioning to assess the contributions of SNP classes defined by functional annotations. Across 11 diseases they concluded that the majority of 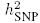 was explained by DHS, despite these containing less than 20% of all SNPs. For Figure 5, we perform a similar analysis using the 10 traits we have in common with their study (for 9 of these, we are using the same data). When we copy Gusev *et al.* and assume the GCTA Model with α = –1, we estimate that on average DHS contribute 86% (SD 4) of 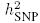, close to the value they reported (79%). When instead we assume the LDAK Model (with α = –0.25), the estimated contribution of DHS reduces to 25% (SD 2). Under the LDAK Model, DHS are predicted to contribute 18% of 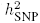 so 25% represents 1.4-fold enrichment. To add context, we also consider “genic” SNPs, which we define as SNPs inside or within 2 kb of an exon (using RefSeq annotations^31^), and “inter-genic,” SNPs further than 125 kb from an exon; these definitions ensure that these two SNP classes are also predicted to contribute 18% of 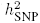 under the LDAK Model. We estimate that genic SNPs contribute 29% (SD 2), while inter-genic SNPs contribute 10% (SD 2), representing 1.6-fold and 0.6-fold enrichment, respectively. When we extend this analysis to all 42 traits, DHS on average contribute 24% (SD 2) of 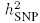, and in contrast to Gusev *et al.*, enrichment remains constant when we reduce SNP density (Supplementary Fig. 14 & 15 and Supplementary Table 8).

Finucane *et al.*^32^ performed a similar analysis, but considered 52 SNP classes and estimated enrichment using LDSC; across nine traits, they identified five classes with >4-fold enrichment, the highest of which, “conserved SNPs,” had 13-fold enrichment. When we use LDAK to estimate enrichment for our 19 GWAS traits, the results are more modest; the highest enrichment is 2.5-fold, with only 1.3-fold enrichment for conserved SNPs (Supplementary Fig. 16).

### Relaxing quality control

For the UCLEB data, we consider nine alternative SNP filterings. Supplementary Figure 17 reports estimates of 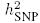 for each trait/filtering, while Figure 6a provides a summary. First we vary the information score threshold: *r_j_* > 0.99, > 0.95, > 0.9, > 0.6, > 0.3 and > 0 (each time continuing to require MAF > 0.01). Simulations suggest that by including all 8.8 M common SNPs (*r_j_* > 0), instead of using just the 353K high-quality ones (*r_j_* > 0.99), we can expect estimates of 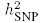 to increase by 50-60% (Supplementary Fig. 18). This is similar to what we observe in practice, as across the 23 traits, estimates of 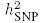 (using α = –0.25) are on average 45% (SD 8) higher. The simulations further predict that, even though the Metabochip provides relatively low coverage of the genome (after quality control, it contains only 60K SNPs, predominately within genes), we can expect estimates of 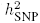 to be approximately 80% as high as those obtained starting from genome-wide genotyping arrays. While we are unable to test this claim directly, it is consistent with our results for height, body mass index and QT Interval, the three traits for which reasonably precise estimates of common SNP 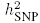 are available^6^ (Figure 6b). For the final three SNP filterings, we vary the MAF threshold: MAF > 0.0025, MAF > 0.001 and MAF > 0.0005 (all with *r_j_* > 0). Across the 23 traits, we find that rare SNPs contribute substantially to 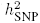: for example, when we use the 17.3 M SNPs with MAF > 0.0005, estimates of 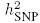 (using α = –0.25 and MAF partitioning) are on average 29% (SD 12) higher than those based on the 8.8M common SNPs (median increase 22%), with rare SNPs contributing on average 33% (SD 5) of 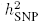 (Figure 6a).

## Discussion

With estimates of 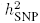 so widely reported, it is easy to forget that calculating the variance explained by large numbers of SNPs is a challenging problem. To avoid over-fitting, it is necessary to make strong prior assumptions about SNP effect sizes, but different assumptions can lead to substantially different estimates of 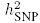. Previous attempts to assess the validity of assumptions have used simulation studies,^14,15^ but this approach will tend to favor assumptions similar to those used to generate the phenotypes. Instead, we have compared different heritability models empirically, by examining how well they fit real datasets.

We begun by investigating the relationship between heritability and MAF. Across 42 traits, we found that best fit was achieved by setting α = –0.25 in (1), which implies that average heritability varies with [MAF(1–MAF)]^0.75^. As explained in Online Methods, the value of α corresponds to the scaling of genotypes. Therefore, our result indicates that the performance (i.e., detection power and/or prediction accuracy) of many penalized and Bayesian regression methods, for example, the Lasso, ridge regression and Bayes A,^33–35^ could be improved simply by changing how genotypes are scaled. Although we recommend α = –0.25 as the default value, with sufficient data available, it should be possible to estimate α on a trait-by-trait basis, or to investigate more complex relationships between heritability and MAF. In particular, with a better understanding of the relationship between heritability and MAF for low frequencies, it may no longer be necessary to partition by MAF when rare SNPs are included.

We also examined the relationship between heritability and LD. To date, most estimates of 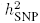 have been based on the GCTA Model; this model can be motivated by a belief that each SNP is expected to have the same effect on the phenotype, from which it follows that the expected heritability of a region should depend on the number of SNPs it contains. By contrast, the LDAK Model views highly-correlated SNPs as tagging the same underlying variant, and therefore believes that the expected heritability of a region should vary according to the total amount of distinct genetic variation it contains. Across our traits, we found that the relationship between heritability and LD specified by the LDAK Model consistently provides a better description of reality.

This finding has important consequences for complex trait genetics. Firstly, it implies that for many traits, common SNPs explain considerably more phenotypic variance than previously reported, which represents a significant advance in the search for missing heritability.^2^ It also impacts on a large number of closely-related methods. For example, LDSC,^10^ like GCTA, assumes that heritability contributions are independent of LD and therefore it also tends to under-estimate 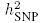. Similarly, we have shown that estimates of the relative importance of SNP classes via SNP partitioning can be misleading when the GCTA Model is assumed.^7,32^ Further afield, most software for mixed model association analyses (e.g., FAST-LMM, GEMMA and MLM-LOCO) use an extension of the GCTA Model,^36–38^ and likewise most bivariate analyses, including those performed by LDSC.^8, 39, 40^ It remains to be seen how much these methods would be affected if they employed more realistic heritability models.

Attempts have been made to improve the accuracy of heritability models via SNP partitioning.^14,15,41^ We find that partitioning by MAF can be advantageous, as it guards against misspecification of the relationship between heritability and MAF when rare variants are included. Figure 3a and Supplementary Figure 7 indicate that the realism of the GCTA Model can be improved by partitioning based on LD; for example, across the GWAS traits, estimates from GCTA-LDMS are on average 16% (SD 2) higher than those from GCTA, and now only 23% (SD 2) lower than those from LDAK. The improvement arises because model misspecification is reduced by allowing SNPs in lower-LD tranches to have higher average heritability. However, Supplementary Table 9 illustrates why we consider such an approach sub-optimal; in particular, SNP partitioning can be computationally expensive, and even with LD-partitioning, model fit tends to be worse than that from LDAK.

While we have investigated the role of MAF, LD and genotype certainty, there remain other factors on which heritability could depend, in particular the available functional annotations of genomes.^42^ For example, our comparison of genic and inter-genic SNPs indicates that the effect-size prior distribution could be improved by taking into account proximity to coding regions. By way of demonstration, Supplementary Table 10 shows that model fit is improved by assuming 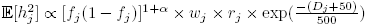, where *D*_*j*_ is the distance (in kb) between SNP *j* and the nearest exon (under this model, genic SNPs are expected to have about twice the heritability of inter-genic SNPs). In general, we believe that modifications of this type will have a relatively small impact; we note that across the 19 GWAS traits, scaling by exp 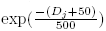 increases model log likelihood by on average only 1.5, much less than the average increase obtained by using α = –0.25 instead of α = –1 (8.9), or by choosing the LD-model specified by LDAK instead of GCTA (17.7), and does not significantly change estimates of 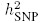. However, with sufficient data, it may be possible to obtain more substantial improvement by tailoring model assumptions to individual traits.

When estimating 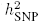, care should be taken to avoid possible sources of confounding. Previously, we advocated a test for inflation of 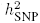 due to population structure and familial relatedness.^3^ The conclusions of a recent paper claiming that 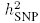 estimates are unreliable,^43^ would have changed substantially had this test been applied (Supplementary Fig. 19). We also recommend testing for inflation due to genotyping errors, particularly before including lower-quality and/or rare SNPs. For the 23 UCLEB traits, we showed that including poorly-imputed SNPs resulted in significantly higher estimates of 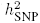, and made it possible to capture the majority of genome-wide heritability despite the very sparse genotyping provided by the Metabochip. We found that including rare SNPs also led to significantly higher 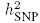. Although sample size prevented us from obtaining precise estimates of 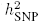 for individual traits, our analyses indicated that for larger datasets, including rare SNPs will be both practical and fruitful in the search for the remaining missing heritability.^2^

## Acknowledgments

Access to Wellcome Trust Case Control Consortium data was authorized as work related to the project “Genome wide association study of susceptibility and clinical phenotypes in epilepsy,” while access to Children’s Hospital of Philadelphia (CHOP) data was granted under Project 49228-1, “Assumptions underlying estimates of SNP Heritability.” We thank Anne Molloy, James Mills and Lawrence Brody for permission to use genotype data from the Trinity College Dublin Student Study.^44^ and Sarah Langley for help accessing the CHOP data. This work is funded by the UK Medical Research Council under grant MR/L012561/1, by the British Heart Foundation under grant RG/10/12/28456, and supported by researchers at the National Institute for Health Research (NIHR) University College London Hospitals Biomedical Research Centre. NC is an ESPOD Fellow from the European Molecular Biology Laboratory, European Bioinformatics Institute, and Wellcome Trust Sanger Institute. SN is a Wellcome Trust Senior Research Fellow in Basic Biomedical Science and is also supported by the NIHR Cambridge Biomedical Research Centre. Analyses were performed with the use of the UCL Computer Science Cluster and the help of the CS Technical Support Group, as well as the use of the UCL Legion High Performance Computing Facility (Legion@UCL) and associated support services.

## Author Contributions

DS and NC performed the analyses. DS and DJB wrote the manuscript with assistance from NC, MRJ, SN and members of the UCLEB Consortium.

## Online Methods

Supplementary Note 1 summarizes the different analyses we performed, and the conclusions we drew from each, while Supplementary Protocol 1 provides step-by-step instructions for estimating 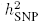 starting from raw genotype data. In general, we assume there are *n* individuals, recorded for *p* covariates and genotyped (either directly or via imputation) for *m* SNPs: the length-*n* vector ***Y*** contains phenotypic values, the *n* × *p* matrix ***Z*** contains covariates, while the *n* × *m* matrix ***S*** contains (expected) allele counts.

### Information score r_j_

Let the vector ***S_j_*** = (*S*_1, *j*_,…, *S_n,j_*)^*T*^ ∈ [0, 2]^*n*^, denote the allele counts for SNP *j* (i.e., *S_j_* is Column *j* of ***S***). Our information score *r_j_* estimates the squared correlation between ***S_j_*** and ***G_j_*** = (*G*_1, *j*_,…, *G_n,j_*)^*T*^ ∈ {0, 1, 2}^*n*^, the true genotypes for SNP *j*. When using imputed data, ***G_j_*** is typically not known; instead for each individual we have a triplet of state probabilities (*p*_*i,j*,0_, *p*_*i,j*,1_, *p*_*i,j*,2_), where 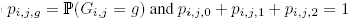. Therefore, we define *r_j_* by taking expectations over the 3^*n*^ possible realizations of *G_j_*.

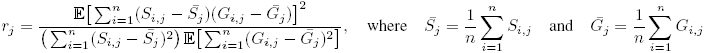

*S_j_* is known, so computing 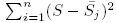 is straightforward. The two expectations can also be calculated explicitly:

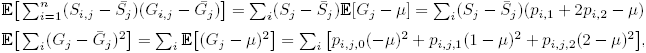

where 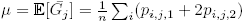. For our analyses, we use expected allele counts (dosages), so 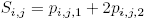. In this case, 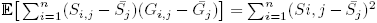 and so the score reduces to 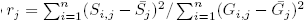. For a directly genotyped SNP, each triplet of state probabilities will be (1,0,0), (0,1,0) or (0,0,1), which will result in *S_i,j_* = *G_i,j_* for all *i* and *r_j_* = 1; so for these, in place of *r_j_*, we use the metric `r2_type0` reported by IMPUTE2.^26^ Additional details on our information score are provided in Supplementary Figure 20.

### Estimating 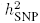

We first construct the *n* × *m* genotype matrix ***X***, by centering and scaling the allele counts for each SNP according to 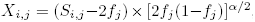, where 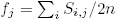. If *w_j_* and *r_j_* denote the LD weight^9^ and information score for SNP *j*, then the LDAK Model for estimating SNP heritability 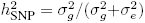 is:

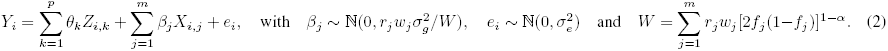

θ_*k*_ denotes the fixed-effect coefficient for the *k*th covariate, β_*j*_ and *e_i_* are random-effects indicating the effect size of SNP *j* and the noise component for Individual *i*, while 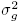 and 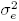 are interpreted as genetic and environmental variances, respectively. Note that the introduction of *r_j_* is an addition to the model we proposed in 2012.^9^ Model (2) is equivalent to assuming:^45,46^

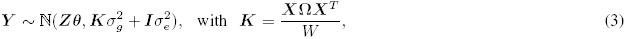

where ***I*** is an *n* × *n* identity matrix and Ω denotes a diagonal matrix with diagonal entries (*r*_1_*w*_1_,…, *r*_*m*_*w*_*m*_). The kinship matrix ***K***, also referred to as a genetic relationship matrix (GRM)^1^ or genomic similarity matrix (GSM),^47^ consists of average allelic correlations across the SNPs (adjusted for LD and genotype certainty). Model (3) is typically solved using REstricted Maximum Likelihood (REML), which returns estimates of θ_1_,…, θ_*p*_, 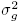 and 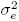.^12^

The heritability of SNP *j* can be estimated by 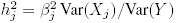, which under Model (2), and assuming Hardy-Weinberg Equilibrium,^48,49^ has expectation

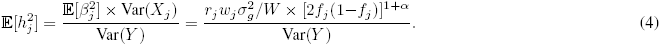

If *P*_1_ and *P*_2_ index two sets of SNPs of size ∣*P*_1_∣ and ∣*P*_2_∣, then under the LDAK Model, they are expected to contribute heritability in the ratio *W*_1_: *W*_2_, where 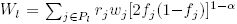. The GCTA Model corresponds to setting *w_j_* = *r_j_* = 1, in which case 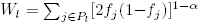. Most applications of GCTA have further assumed α = –1, so that *W_l_* = ∣*P*_l_∣, which corresponds to the assumption that SNP sets are expected to contribute heritability proportional to the number of SNPs they contain.

Model (2) assumes that all effect-sizes can be described by a single prior distribution. This assumption is relaxed by SNP partitioning. Suppose that the SNPs are divided into tranches *P*_1_,…, *P_L_* of sizes ∣*P*_1_∣,…, ∣*P_L_*∣; typically these will partition the genome, so that each SNP appears in exactly one tranche and _*l*_∣*P_l_*∣ = *m*, but this is not required. This correspond to generalizing Model (2), so that SNPs in Tranche l have effect-size prior distribution 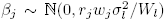. Letting 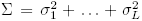, then 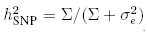, while 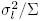 represents the contribution to 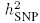 of SNPs in Tranche l. This model can equivalently be expressed as 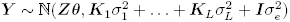, where ***K**_l_* represents allele correlations across the SNPs in Tranche *l*.

For analyses under the LDAK Model, we used LDAK v.5; for analyses under the GCTA Model, we used GCTA v.1.26. For about a third of GCTA-LDMS analyses, the GCTA REML solver failed with the error “information matrix is not invertible,” in which case we rerun using LDAK (while the GCTA and LDAK solvers are both based on Average Information REML,^30,50^ subtle differences mean that when using a large number of tranches, one might complete while the other fails). For the few occasions when both solvers failed, we instead used “GCTA-LD” (i.e., SNPs divided only by LD, rather than by LD and MAF), which we found gave very similar results to GCTA-LDMS for traits where both completed (Supplementary Fig. 7). For diseases, we converted estimates of 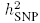 to the liability scale based on the observed case-control ratio and assumed prevalence.^28,29^ In general, we copied the prevalences used by previous studies; however for tuberculosis, where no previous estimate of 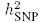 is available, we derived an estimate of prevalence from World Health Organization data^51^ (Supplementary Note 2).

### LDSC

Originally designed as a way to quantify confounding in a GWAS, LDSC^10^ also provides a method for estimating 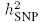, which requires only summary statistics from single-SNP analysis (rather than raw genotype and phenotype data). LDSC is based on the principal that in a single-SNP analysis, the χ^2^(1) test statistic for SNP *j* has expected value 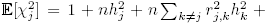*na_j_*, where 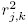 denotes the squared correlation between SNPs *j* and *k*, while *a_j_* represents bias due to confounding factors (e.g., population structure and familial relatedness).^10^ Under a polygenic model where every SNP is expected to contribute equally (i.e., 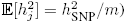), and the (widely-used) assumption that the bias is constant across SNPs (*a_j_* = *a*), we have 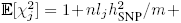*na*, where 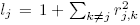 is referred to as the LD Score of SNP *j* (as it is not feasible to compute pairwise correlations across all SNPs, in practice these are approximated using a sliding window of, say, 1 centiMorgan). Therefore, LDSC estimates 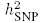 and *a* by regressing test statistics on LD Scores. In the absence of confounding (*a* = 0), LDSC can be viewed as estimating 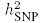 under the GCTA Model with α = –1 (as this satisfies the assumption that every SNP is expected to contribute equal heritability). As the authors of LDSC point out,^10^ it is straightforward to accommodate alternative relationships between 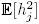 and MAF (i.e., α ≠ –1) by changing how genotypes are scaled when computing LD Scores, and potentially genotype certainty could be accommodated. However, the similarity with the GCTA Model appears intrinsic to LDSC; while the assumption that heritability is independent of LD can be relaxed via SNP partitioning,^41^ we can not envisage how the method could be modified to accommodate the LDAK SNP weights. For LDSC analyses, we used LDSC v.1.0.0 both for calculating LD Scores and estimating 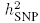.

### Accommodating very large effect loci

Equation (2) assumes that all SNP effect sizes can be modeled by a single Gaussian distribution. Estimates are generally robust to violations of this assumption,^9^ but problems can occur when individual SNPs have very large effect sizes, because a single Gaussian distribution cannot accommodate both these SNPs and the very many with small effect sizes. This is a common concern when analyzing autoimmune traits for which the major histocompatibility complex (MHC) can contribute substantial heritability. In response to this problem, some authors exclude MHC SNPs from analyses.^7,30,52,53^ Another approach is to model effect sizes as a mixture of Gaussians,^35,54^ but this is not computationally feasible for millions of SNPs and many thousands of individuals. Therefore, our proposed strategy is to first identify SNPs with *P* < 10^−20^ from single-SNP analysis, to prune these using a correlation squared threshold of 0.5, then to include those which remain as fixed-effect covariates. Thus in place of Equation (3), we assume 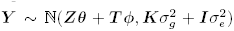, where columns of the matrix ***T*** contain allele counts of the highly-associated SNPs (i.e., ***T*** is a submatrix of ***S***), and the vector φ represents their effect sizes. In contrast to standard (non-SNP) covariates, the variance explained by ***T*** counts towards SNP heritability: 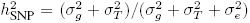, where 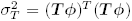 Supplementary Figures 21 & 22 provides further details. In particular, we appreciate that our definition of highly-associated is somewhat arbitrary, so we confirm that estimates of 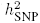 are almost unchanged if instead we use *P* < 5 × 10^−8^.

### Datasets and phenotypes

When searching for GWAS datasets, we preferred those with sample size at least 4 000 to ensure reasonable precision of 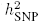.^55^ In total, our datasets were constructed from 40 independent cohorts, all of which have been previously described (see Supplementary Tables 11 & 12 for references and details of how cohorts were merged to form datasets). For the UCLEB data, there were in total 28 quantitative traits with measurements recorded for at 7 000 individuals. For each of these, we quantile normalized, then applied a test for inflation due to genotyping errors (Supplementary Fig. 13). Specifically, our test, inspired by Bhatia *et al.*^56^ and valid for quantitative phenotypes where individuals are recruited from multiple cohorts, first estimates 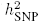 using only pairs of individuals in different cohorts, then using only pairs of individuals in the same cohort; a significant difference between the two estimates indicates possible inflation due to genotyping errors. We excluded five traits that showed evidence of inflation (*P* < 0.05/28), leaving us with 23: height, weight, body mass index, waist circumference, forced vital capacity, one second forced vital capacity, systolic blood pressure (adjusted), diastolic blood pressure (adjusted), PR Interval, QT Interval, Corrected QT Interval, QRS Voltage Product, Sokolow Lyon, glucose, insulin, total cholesterol (adjusted), LDL cholesterol (adjusted), triglyceride (adjusted), viscosity, fibrinogen, Interleukin 6, C-reactive Protein and haemoglobin. Approximately 40% of individuals were receiving medication to reduce blood pressure, 25% to reduce lipid levels, so where indicated, phenotypes had been adjusted for this: for individuals on medication, their raw measurements had been increased either by adding on (blood pressure) or scaling by (lipid levels) a constant.^57,58^ We note that some pairs of traits are highly correlated. However, as the overall correlation is not that extreme (we estimate the effective number of independent traits to be about 15), and most of our UCLEB analyses serve to support conclusions drawn from the GWAS traits, we decide to retain all 23 traits (rather than, say, consider only a subset). See Supplementary Note 3 for further details on phenotyping.

### Quality control

We processed each of the 40 cohorts in identical fashion; see Supplementary Note 4 for full details. In summary, after excluding apparent population outliers, samples with extreme missingness or heterozygosity, and SNPs with MAF < 0.01, call-rate < 0.95 or *P* < 10^−6^ from a test for Hardy-Weinberg Equilibrium, we imputed using IMPUTE2^26^ and the 1000 Genome Phase 3 (2014) Reference Panel.^27^ When merging cohorts to construct the GWAS datasets, we retained only autosomal SNPs which in all cohorts have MAF > 0.01 and *r_j_* > 0.99 (using IMPUTE2 `r2_type2` in place of *r_j_* for directly genotyped SNPs). For the 8 UCLEB cohorts, we applied these filters only after merging. We only relax quality control for the analyses of the UCLEB data where we explicitly examine the consequences of including lower-quality and rare SNPs. When possible, the matrix ***S*** contains expected allele counts (dosages); i.e., *S*_*i,j*_ = *P*_*i,j*,1_ + 2 × *p*_*i,j*,1_, where *p*_*i,j*,1_ and *p*_*i,j*,2_ denote the probabilities of allele counts 1 and 2, respectively. If hard genotypes are required, for example when using LDSC to compute LD Scores,^10^ we round *S_i,j_* to the nearest integer. As this was only necessary when considering high-quality SNPs (*r_j_* > 0.99), we expect this rounding to have negligible impact on results. For each trait, Table 1 reports m, the total number of SNPs after imputation, and 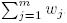, the sum of SNP weights; the aim of these weights is to remove duplication of signal due to LD and their sum can loosely be interpreted as an effective number of independent SNPs. For the GWAS datasets, *w_j_* ranges from 79K to 125K. By contrast, when restricted to only high-quality SNPs, the UCLEB data has *w_j_* = 39 K, reflecting that the Metabochip directly captures a much smaller amount of genetic variation than standard genome-wide SNP arrays.

When analyzing quantitative traits, genotyping errors will tend only to be a concern when there are systematic differences between phenotypes across cohorts, and this is something we are able to explicitly test (Supplementary Fig. 13). However, for disease traits, when cases and controls have been genotyped separately (as is the design of most of our GWAS datasets), any errors will almost certainly correlate with phenotype and therefore cause inflation of 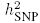.^9,29^ To test the effectiveness of our quality control for the GWAS traits, we construct a pseudo case-control study using two control cohorts; we confirm that the resulting estimate of 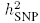 is not significantly greater than zero, suggesting that the quality control steps we use for the GWAS datasets are sufficiently strict (Supplementary Note 5).

Accurate estimation of 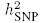 requires samples of unrelated individuals with similar ancestry. Prior to imputation, we removed ethnic outliers identified through principal component analyses (Supplementary Fig. 23). Post imputation, we computed (unweighted) allelic correlations using a pruned set of SNPs, then filtered individuals so that no pair remained with correlation greater than *c*, where –*c* is the smallest observed pairwise correlation (*c* ranges from 0.029 to 0.038, depending on dataset). For our datasets, this filtering excluded relatively few individuals (on average 3.8%, with maximum 11.6%). For all analyses, we include a minimum of 30 covariates: the top 20 eigenvectors from the allelic correlation matrix just described, and projections onto the top 10 principal components computed from 1000 Genomes samples.^27^ For the 19 GWAS traits, we also include sex as a covariate, while for intraocular pressure and wide range achievement test scores, we additionally include age. Supplementary Figure 24 reports the proportion of phenotypic variance explained by each covariate. To check our filtering and covariate choices, we estimate the inflation of 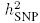 due to population structure and residual relatedness^3^ (Supplementary Fig. 19). For the GWAS traits, we estimate that on average 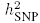 estimates are inflated by at most 3.1%, with the highest observed for ischaemic stroke (7.1%). For the 23 UCLEB traits, the average inflation is 0.3% (highest 2.3%).

### Single-SNP analysis

Supplementary Figure 25 provides Manhattan Plots from logistic (case-control traits) and linear regression (quantitative traits), performed using PLINK v.1.9. These analyses provide the summary statistics required by LDSC. For the GWAS traits, we identified highly-associated SNPs (*P* < 10^−20^) within the MHC for 6 of the GWAS traits (rheumatoid arthritis, type 1 diabetes, psoriasis, ulcerative colitis, celiac disease and multiple sclerosis), while rs2476601, a SNP within *PTPN22*, is highly associated with both rheumatoid arthritis and type 1 diabetes.^59,60^ For the UCLEB traits, we find highly associated SNPs within *SCN10A* (PR Interval), *APOE* (total cholesterol, LDL cholesterol and C-reactive protein) and *ZPR1* (triglyceride levels). For heritability analysis, these SNPs were pruned, then included as additional fixed-effect covariates as described above.

### Computational requirements

The most time-consuming aspect of analysis was genotype imputation; for a typically-sized cohort (∼3 000 individuals) this took approximately one CPU-year (i.e., a few days on a 100-node cluster). Next is computation of SNP weights, which for the GWAS traits (∼4M SNPs) took approximately one CPU-month (again, this can be near-perfectly parallelized). Finally, solving the mixed-model via REML would take between a few minutes for the smaller traits (∼5 000 individuals) and a few hours for the largest (∼ 14 000 individuals). Memory-wise, the most onerous task is solving the mixed-model, for which memory demands scale with *n*^2^; however, even for the largest dataset, this was less than 5 Gb (when using multiple kinship matrices, LDAK allows for these to be read on-the-fly, so that the memory demands are no higher than when using only one).

